# Analysis of noisy transient signals based on Gaussian process regression

**DOI:** 10.1101/2022.09.27.509665

**Authors:** I. Baglaeva, B. Iaparov, I. Zahradník, A. Zahradníková

## Abstract

Dynamic systems such as cells or tissues generate, either spontaneously or in response to stimuli, transient signals that carry information about the system. Characterization of recorded transients is often hampered by a low signal-to-noise ratio (SNR). Reduction of the noise by filtering has limited use due to partial signal distortion. Occasionally, transients can be approximated by a mathematical function, but such a function may not hold correctly if recording conditions change. We introduce here the model-independent approximation method for general noisy transient signals based on the Gaussian process regression (GPR). The method was implemented in the software TransientAnalyzer, which detects transients in a record, finds their best approximation by the Gaussian process, constructs a surrogate spline function, and estimates specified signal parameters. The method and software were tested on a cellular model of the calcium concentration transient corrupted by various SNR levels and recorded at a low sampling frequency. Statistical analysis of the model data sets provided the error of estimation <7.5% and the coefficient of variation of estimates <17% for peak SNR=5. The performance of GPR on signals of diverse experimental origin was even better than fitting by a function. The software and its description are available on GitHub.

**Statement of Significance:** Transient signals convey information on the state and function of the studied system. However, the estimation of their characteristic parameters is complicated by the noise present in the recordings. Methods used for noise reduction have various disadvantages, such as distortion of the time course by filtering, the difficult superposition of many transients for accurate averaging, or a lack of a model for data fitting. In this work, we present a general method for the automatic analysis of noisy transient signals based on Gaussian process regression and its implementation in Python. The method can analyze recorded transients reliably at peak SNR ≥ 2 with a precision equivalent to the model-fitting methods.

## Introduction

Transient signals regulate many processes in biology. Their study is important for the understanding of the underlying mechanisms. These signals are characterized by specific parameters, for instance, the peak amplitude, the time to peak, the duration at half maximum, or the rise and decay times, which can be read from the recorded traces if the signal-to-noise ratio is sufficiently large and the characteristics of interest are not degraded by the excess noise. In reality, the recorded signals are contaminated by excess noise that has to be suppressed before data analysis.

Currently available software products for transient signal analysis have various limitations. The commercial software either works only with specific hardware (1) or requires extensive manual input (2). Open source software that provides detection and analysis of transient signals is intended for a very specific type of signal such as cardiac calcium sparks or spikes (3–7) or neuronal calcium spikes (8) and/or is written for a proprietary software platform (3, 9). The methods of signal detection range from manual specifications of baseline and transient intervals (2) through matrix-based image analysis (6, 7) to solving optimization problems by gradient methods, maximal likelihood, and/or deep learning (3, 4, 8). The analysis of detected signals proceeds by direct reading of values at predefined temporal and/or amplitude positions (model-independent; (2)), fitting of phenomenological or model-based functions (3, 10, 11), or averaging (2). Algorithms that do not use a data fitting function use linear interpolation between recorded data points to estimate a parameter value between recorded data points and thus may report values with errors proportional to the local instances of signal distortion. In practice, the peak signal-to-noise ratio (SNR), measured as the ratio of the peak transient amplitude to the noise level, may be below 10 (4, 5, 11), that is, much smaller than acceptable for analysis by such simple algorithms.

Several noise-suppression techniques are generally applied. The signal averaging method is applicable only if the generated transients represent stationary signals and have a well-defined onset time or timing by activating stimuli to synchronize their superposition. As methods of choice, a band-pass filter (12) or approximation by a known function (10, 11) were used with non-stationary transients. Application of a filter to the data record may be of some help, but often at the cost of some degradation of the signal time course inherent to any filter. The approximation of transients by a specific mathematical function depends on the quality of the mathematical model. Less exact or phenomenological models may introduce systematic errors. On the other hand, the application of a mathematical model of a transient with a complex structure, or analysis of signals recorded under unstable conditions, may need certain corrections in the parameter setting or in the model itself, disqualifying them from the automatic analysis.

In general, the shape of the transient can be very complex and variable. For the approximation of a general transient, we considered the Gaussian Process Regression (GPR) method (13), originally developed to approximate a multidimensional dataset obtained from measurements of complex structures in a variety of fields of sciences such as mining, engineering, agriculture (14), epidemiology (15) and cardiovascular modeling (16). It also found applications in solving other problems such as image denoising (17) and signal separation (18). In the approximation problem of a one-dimensional dataset, we turned to the GPR as a nonparametric regression model, in which the model structure of a transient is not predetermined, e.g., by logistic or polynomial functions, but is constructed as the probability distribution over the functions defined by the kernel and conditioned by the given data set (13). In this work, we describe and test an automatic algorithm and software for the analysis of individual noisy transient signals based on nonparametric regression that allows the user to approximate transients with a different steepness of the rising and decaying part of their time course. We demonstrate the ability of the software to automatically detect calcium transients and estimate their characteristic parameters reproducibly even at the peak SNR as low as 2. The error of estimation was less than 7.5% and the coefficient of variation of estimates was less than 17% for peak SNR ≥ 5. The source code and the executable file on GitHub (https://github.com/IuliiaBaglaeva/TransientAnalyzerUI) listed in Zenodo (19) are freely available.

## Methods

This section describes the methods of transients detection and their approximation by the GPR approach. We also define here parameters used to characterize individual transients and describe the statistical analysis used to evaluate the precision of parameter estimation by TransientAnalyzer.

The software TransientAnalyzer was written in Python version 3.7. The inputs are NumPy (20) arrays of recorded time points and signal amplitudes. The TransientAnalyzer sequentially performs the detection of transients in the noisy records, the best approximation of individual transients with a smooth continuous function, and the calculation of characteristic parameter values of the smoothed transients.

The software TransientAnalyzer is presented and deposited as a Graphical User Interface application (https://github.com/IuliiaBaglaeva/TransientAnalyzerUI/releases) and as a Python module for developers in GitHub (https://github.com/IuliiaBaglaeva/TransientAnalyzer) together with a listing on Zenodo with a Digital Object Identifier (DOI) 10.5281/zenodo.7039241 so that it is uniquely citable.

### Baseline correction

Baseline drift may complicate the analysis of transients. The TransientAnalyzer provides optional baseline correction based on the improved modified polynomial method (imodpoly of 4^th^ order (21)) implemented in the pybaselines library (22).

### Determination of the transients’ polarity

The TransientAnalyzer automatically determines the polarity of the transients, i.e., whether they fall or rise relative to the baseline. The probability density function was constructed from all signal values and approximated by two Gaussians using the Expectation-Maximization (EM) algorithm (23). It is assumed that the Gaussian with the larger weight corresponds to the baseline. Then, if the mean of the component with the larger weight is smaller than the mean of the second component, the transients were positive or rising; otherwise, they were negative or falling. In the forthcoming description of the method, the transients are assumed to be positive. The same method is applied to negative transients after changing the sign of the signal.

### Detection of transients

The detection of a transient in the record is based on finding the rapid rise of the signal using the difference function *D*(*t*) between the recorded noisy original trace (*f*(*t*)) and the smoothed original trace (*f1*(*t*)) calculated by a moving average (boxcar) filter at each time point. For the demonstrated example of calcium transients, the difference *f*(*t*) - *f1*(*t*) shows a sharp maximum at the onset of the transient (Figure 1). The correct number of detected transients and their starting points are influenced by the proper setting of the boxcar and the prominence parameters of the moving average filter. The boxcar size (*w1*) specifies the number of points averaged by the filter and the prominence specifies the threshold of detected peaks relative to the surrounding baseline (24). An improper prominence setting may influence the number of detected transients, especially at a low SNR. An improper boxcar size may affect the estimate of the transients’ start point, depending on the kinetics of the transient and the sampling frequency. The software allows finding the correct number of transients and their approximate start time points even in the case of excess noise by interactive corrections.

**Figure 1.**
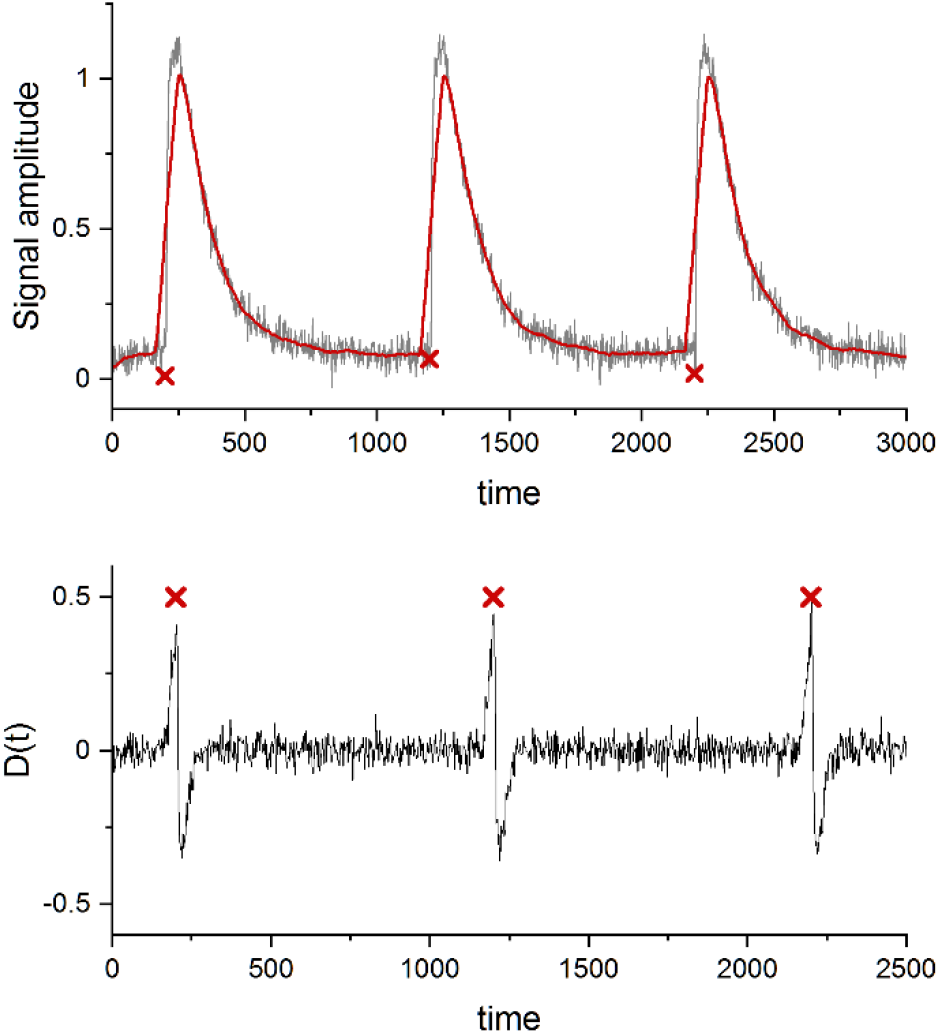
The detection of transients and determination of their start points. Top: The simulated record of transients with a signal-to-noise ratio of 10 (black), and its moving average-filtered trace (red) at a boxcar size of 20 points and prominence of 1. Bottom: The difference *D*(*t*) between the recorded and the filtered traces. Red crosses indicate the times of peak difference and thus the approximate starting times of the transients.

The difference function *D*(*t*) contains the noise of the original signal, hence a large instance of noise can impede the finding of the correct position of its peak. To keep the automatic transient detection functional at a low SNR, the original signal may be smoothed by an additional narrower boxcar filter of size *w2* giving the trace *f2*(*t*). In this case, the *D*(*t*) is calculated as the difference *f2*(*t*) – *f21*(*t*), where *f21*(*t*) is the original signal after being sequentially smoothed by boxcar filters of sizes *w2* and *w1*. The difference between the two smoothed traces has less noise and the difference peak can be always found. In such a case, the estimated start point of the transients would be time-shifted to the right by the rounded-up value of (*w*2 – 1)/2.

The interval between two consecutive start points of the detected transients is used for setting the time interval of the record to be approximated by the GPR algorithm in one run. If the real start values are known, e.g., from the timing of excitation stimuli, the automatic detection step described above may be overridden by supplying the file with a list of stimulus time points, which will demarcate the points of a transient used for approximation by the GPR. The time point of the stimulus can be also used to estimate the delay between the stimulus and the onset of the transient.

### Approximation of the transient by GPR

The Gaussian Process Regression is a type of nonlinear Bayesian regression (13) with the prior distribution being a multivariate Gaussian distribution with an arbitrary mean (usually set to 0) and a total *N*-dimensional covariance matrix ***K***, where the dimension *N* is the number of samples, with the elements of the matrix

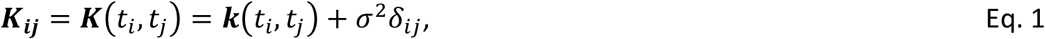

where *t*_i_, *t*_j_ are the points on the time axis and *k*(*t_i_*, *t_j_*) is the covariance matrix of the noiseless signal, *σ* is the noise amplitude of the analyzed signal and *δ_ij_* is the Kronecker delta. In the Gaussian process, the covariance matrix plays a crucial role as it defines the relationship between the points and hence the behavior of the process behind the data. Since the process is not stationary and the analyzed transients may display a positive skewness, the kernel was extended by adding a linear term to consider the fast rise and slow decay of the signal. Mathematically, it is defined as a variant of the Gibbs kernel (25):

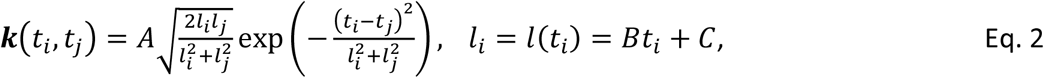

where the parameters *A*, *B*, and *C* define the amplitude and temporal properties of the kernel. The parameters σ (Eq. 1) and *A*, *B*, *C* (Eq. 2) are estimated by maximization of the log marginal likelihood (Eq. 2.30 in ref. (13)) using the limited-memory Broyden, Fletcher, Goldfarb, and Shanno bound (L-BFGS-B) algorithm (26). The GPR model of the transient was built using GPFlow, a Tensorflow-based Python library (27).

To prevent overfitting of the analyzed signal by GPR (i.e., to prevent the approximation function from following the noise) right at the start of the transient (small values of *i* in Eq. 2), the parameter *C* has a lower bound, equal to half of the minimal time step of the signal.

### The characteristics of a transient

The GPR determines the best approximation of the transient at each time point of the original record. It should be noted that the time points may be sampled at irregular intervals. The time resolution of the approximated transients can be increased by finding an appropriate continuous function. To this end, the GPR predictions are sampled at time intervals shorter than those of the input data. In the demonstrated cases, the GPR time interval was set to 100 μs while the recorded sampling period was 2.5 ms. For a further increase in resolution, a surrogate continuous cubic spline model based on the GPR predictions was used for the sake of computational efficiency. This allows the TransientAnalyzer to determine the values of characteristic parameters at increased amplitude and time resolution.

The characteristic parameters of a transient are defined in Figure 2. The start time of the transient origin, *t_0_*, is determined as the intersection between the surrogate spline function of the Gaussian process approximating the analyzed transient, and the baseline estimated as the value of the Gaussian process (GP) describing the preceding transient at its endpoint. For the first transient in the record, the baseline is calculated as the average of the last 80 data points before its starting point. The intersection was found using the Heavy ball method (28), which is the gradient descent algorithm with momentum to speed up convergence. The amplitude is defined as the difference between the peak value of the surrogate spline function describing the transient and the baseline. The intervals between *t*_0_ and points at specific amplitude levels of the rising (*r*_x%_; time to peak (*TTP*) for x = 100%) and decaying parts of the transient (*d*_(100-x)%_) are estimated and used to determine the kinetic characteristics of the transient.

**Figure 2.**
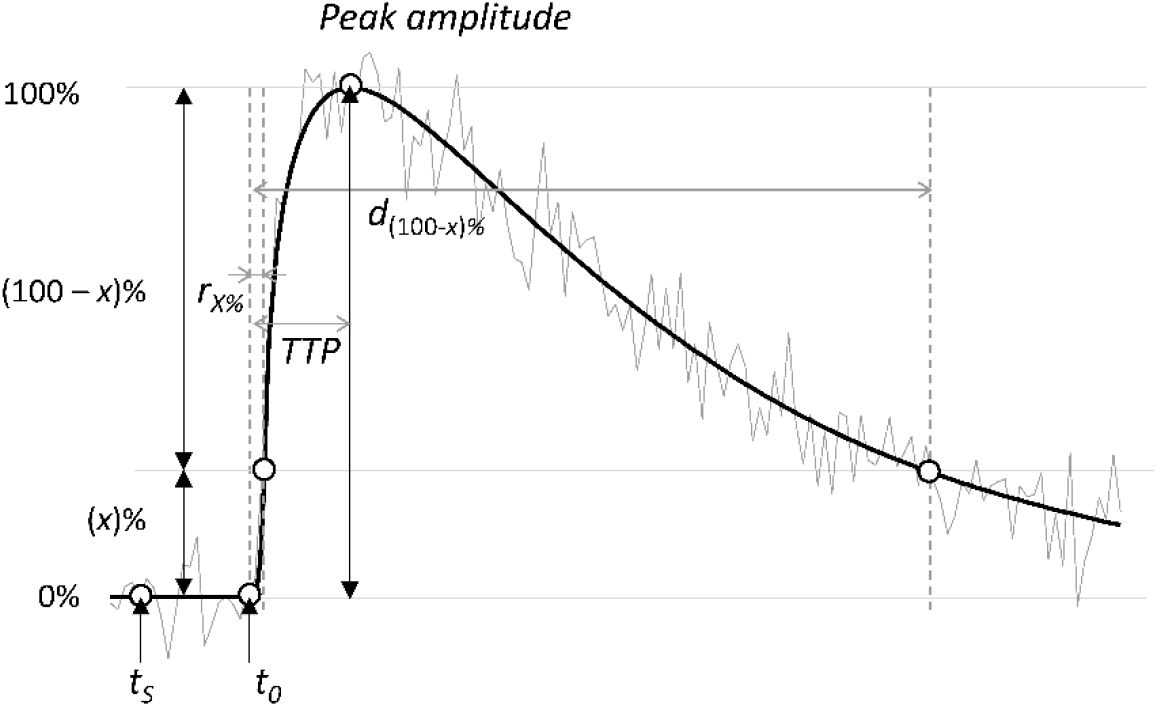
Definition of the characteristic parameters of a transient signal. *t*_S_ - the stimulus time (if a stimulus is present); *t*_0_ - the start time; *r_x%_* – the interval between *t*_0_ and the time to the rise of the signal by *x*% of the amplitude; *TTP* - time to peak; *d*_(100-x)%_ - the interval between *t*_0_ and the time to the decay of the signal from the peak amplitude value by (100-*x*)%. Grey line - an example of a transient signal. Black line - the surrogate spline function for the best GPR approximation of the transient.

Derived parameters that characterize the transient are calculated as described in Table 1. The user may still easily calculate other temporal parameters of interest, such as the time from the peak to a selected level of decay (*d*_x%_ - TTP).

**Table 1.** Secondary kinetic parameters.

| Parameter | Formula |
| --- | --- |
| Delay | $t_0 - t_s$ |
| full duration at half-maximum, FDHM | $d_{50\%} - r_{50\%}$ |
| rise time $t_{x-(100-x)\%}$ | $r_{(100-x)\%} - r_{(x)\%}$ |
| decay time $t_{(100-x)\%-x\%}$ | $d_{(x)\%} - d_{(100-x)\%}$ |
| time to $x\%$ decay from the peak | $d_{(x)\%} - TTP$ |

The kinetic parameters were determined using the bisection method (29), while the peak amplitude, *t*_0_, and time-to-peak (TTP) were determined using the L-BFGS-B method (26).

### Statistics

Statistical analysis was performed in Origin (OriginLab, Ver. 2022b). In the demonstrated example, the signal record was represented by 40,000 points of simulated calcium signal (*t_i_*, *[Ca]_i_*) sampled at 2.5 ms intervals. The time interval of the signal containing one transient approximated by the GPR algorithm was 400 samples (1000 ms). Differences between experimental data and their approximation were quantified as mean squared error.

## Results and Discussion

Generally, the transients are likely to have various kinetics so that their rise time and decay time can differ by an order of magnitude or more. Therefore, any linear time-invariant (LTI) filter, for example, the Savitzky-Golay filter (12), fails to smooth the transient without substantial distortion. It may either overfit the signal and leave a significant noise level if the boxcar size is small or the cutoff frequency is high, or underfit and distort the fast signal dynamics at large boxcar sizes or small cutoff frequencies. In addition, LTI filters significantly shift the onset and the peak time of a transient (compare the black and red line of Figure 1). Such signal distortions may be unacceptable for more precise measurements.

The GPR method with the commonly used stationary kernels such as Squared Exponential or Matern kernels (13) would also fail to approximate the complex transients correctly, similarly to LTI filters, because of the use of a common length scale for the whole transient. We solved this problem by increasing the length scale linearly with the progressing time in the Gibbs kernel (the linear term in Eq. 2) and thus account for the changing dynamics of the transient. This kernel extension allows a better approximation of both, the short and the long time scale processes present in transient signals.

The performance of the TransientAnalyzer was tested on simulated calcium transients generated by a model of human cardiomyocyte electrophysiology (30), which solved the cytosolic calcium concentration response to the incoming action potential. The simulations were conducted in the OpenCOR software (31) using the CVODE numerical solver with a time step of 0.01 ms and recorded at a sampling frequency of 400 Hz. To assure that the estimated parameters have a consistent coefficient of variation, a sequence of 100 identical calcium transients, separated by 1 000 ms start-to-start, was generated and combined with Gaussian noise. The noise was simulated using NumPy (20) to yield an SNR at the peak of the transient in the range of 2 to 100. The noise level σ was defined as

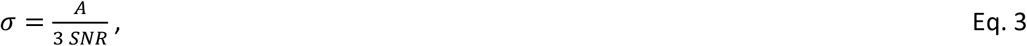

where *A* is the peak amplitude of the simulated transient, estimated as the difference between the signal at the peak and the baseline value.

The TransientAnalyzer could approximate the detected transients well even at very low SNR values; nevertheless, irregular artifacts could be present in the approximated traces especially when the signal approached the baseline (Figure 3). Since such irregularities were not present in the simulated transients, they had to arise from the slow components of the random noise. In experiments, the actual time course of the signal may not be *a priori* inferred; therefore, such distortions have to be accepted as a true signal variation in a model-free analysis.

**Figure 3.**
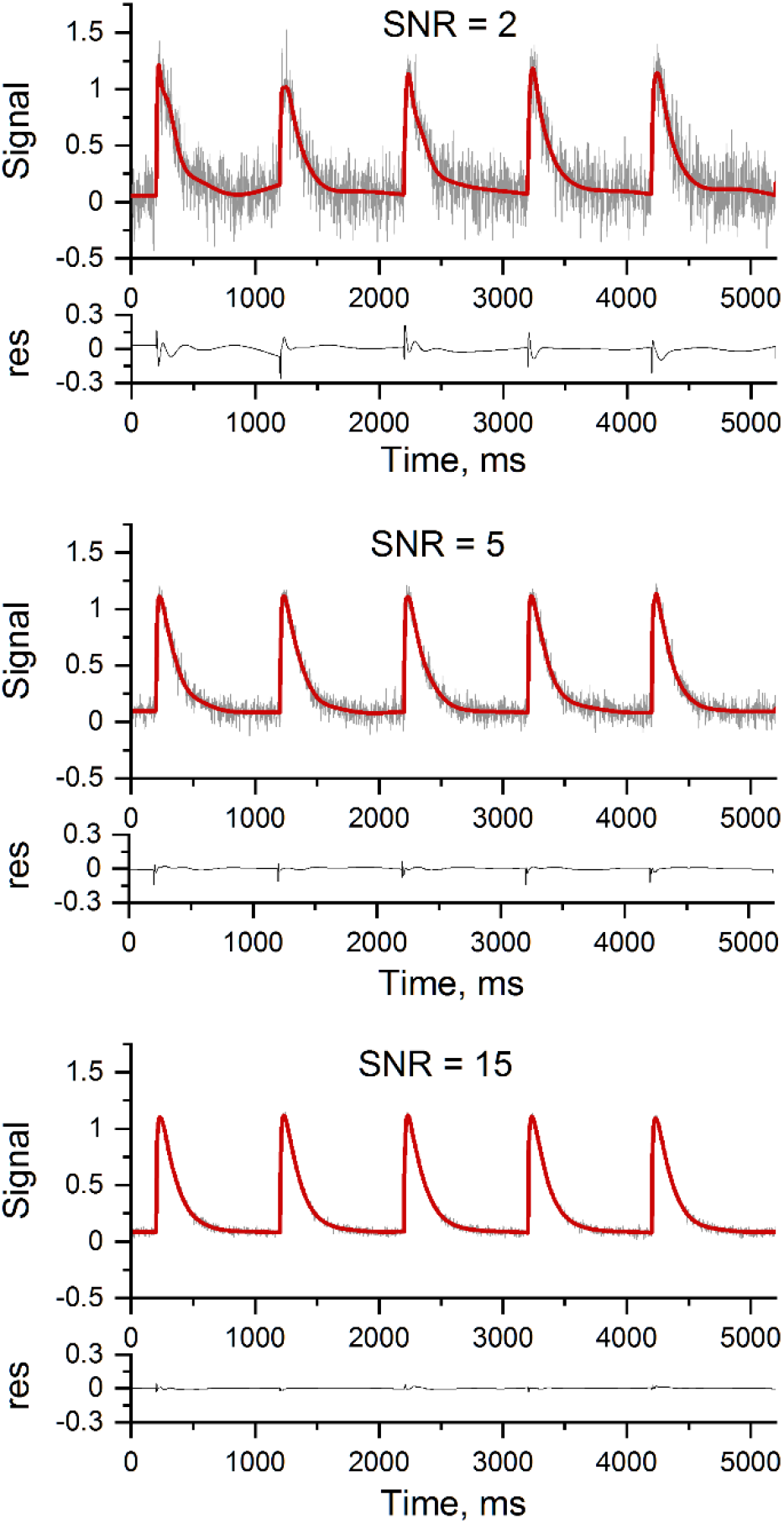
Approximations of transients by the Gaussian process regression. Signal (a.u.) - the examples of simulated noisy transient signals at the indicated peak SNR values with the best approximation curves (red lines); res - residuals.

The parameter values estimated for individual transients were compared with their true values, calculated from the corresponding theoretical transients, by performing statistics on 100 noisy transients for each peak SNR value (Figure 4). At low peak SNR values (< 5), the detection algorithm correctly identified all transient events at the optimized parameter settings of the filter. However, systematic errors were present in the kinetic parameters characterizing the rising part of the transient (Figure 4, left panel). Although at peak SNR = 2 the systematic error was up to 25% in the case of *t*_10-90_%, this amounted to less than two samples. The average values of the remaining estimated parameters were within 2.5% of the true values at all peak SNRs.

**Figure 4.**
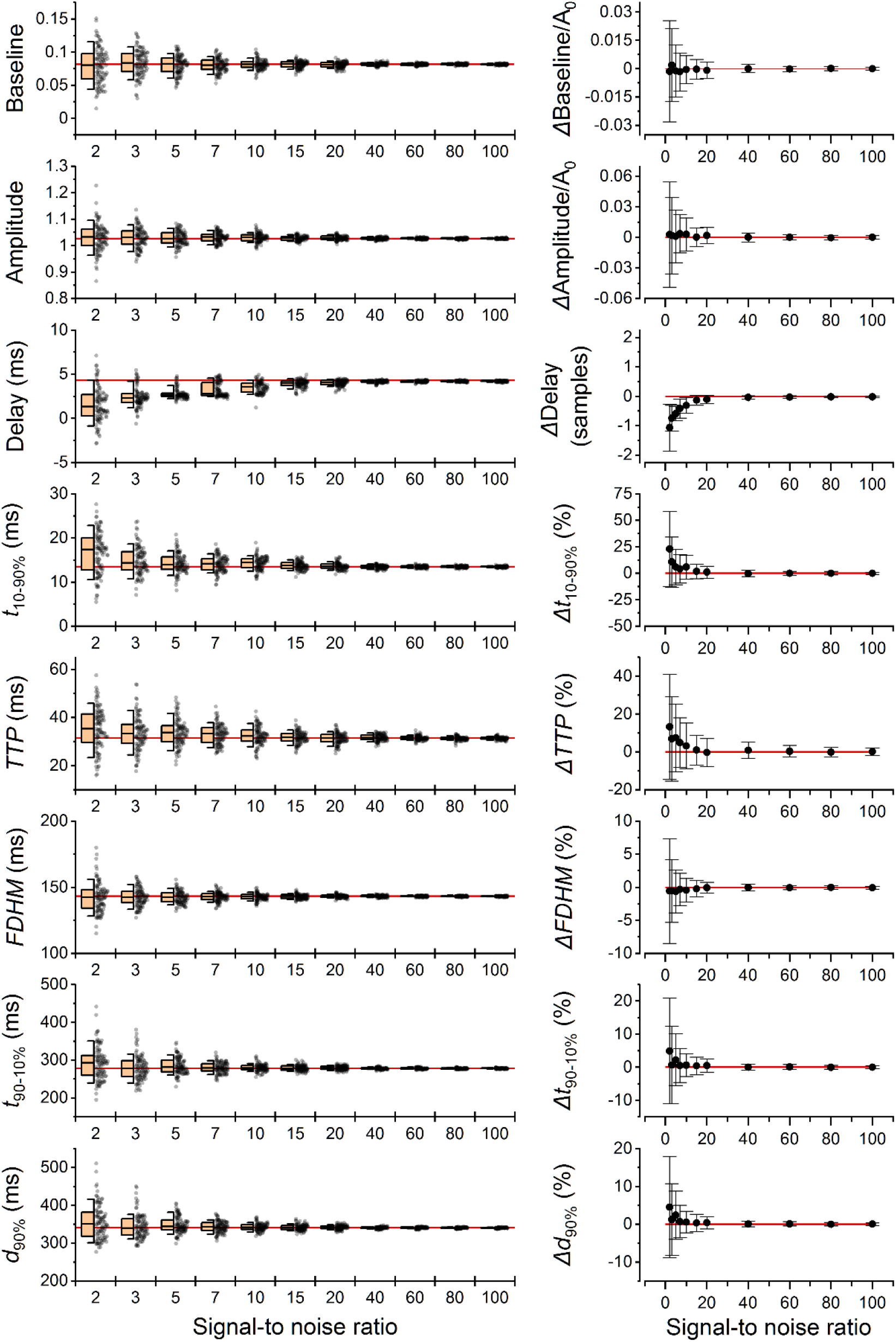
Statistical analysis of estimates of the characteristic parameters of transients by the GPR method. Left: Distribution of the parameter estimates for 100 individual transients at different peak SNR values. The parameter estimates (gray dots) are shown together with the half-box statistic (left to the data). The black line inside the half-box represents the median, whiskers show the 10% and 90% percentile, and the half-box shows the 25% and 75% percentile. The red lines indicate values estimated for the theoretical model of the transient. Right: Systematic errors (circles) of the parameter estimates, calculated as the difference between the mean of estimates at the given peak SNR and the respective parameter value of the theoretical model. The error bars denote standard deviations of estimates. The scale of the Y axis is in units of amplitude for the amplitude and baseline, in units of sample time (2.5 ms) for the delay, and in percentages for the remaining parameters.

The precision of baseline and amplitude estimates was very good for all peak SNR levels, specifically, with a standard deviation of less than 5 % in amplitude units.

The standard deviation of the delay was within one sample time for all peak SNR values; at the peak SNR ≥ 10 it was even below 25% of the sample time (0.6 ms). The largest precision was achieved in the parameter full duration at half-maximum (FDHM) with the systematic error and the coefficient of variation under 1% at all peak SNR values. The largest error was observed in the estimates of the times on the rising part of the transients. For the peak and the *t*_10-90_% rise time, the coefficients of variation were over 25% at peak SNR = 2; this amounted to less than 2 samples. The coefficients of variation of the remaining parameters were less than 7.5% at peak SNR = 5. Taken together, these results demonstrate the robustness of the GPR algorithm and its applicability for the automation of parameter estimation of the transients even in the presence of high noise in the recorded signal.

The data on the accuracy and precision of software for signal analysis at a low signal-to-noise ratio are scarce. For FHDM and Amplitude, the IOCBIO spark detection software reported systematic errors of up to 12% at SNR = 2 (7), while the GPR approach provided less than 1% errors in these parameters (Figure 4). The analysis of precision for IOCBIO was not reported. The software SpikeAnalyzer for analysis of calcium spikes in cardiac myocytes (3), based on the model function derived for the specific time course of calcium release flux (11), reported systematic errors of up to 7% and coefficients of variation of up to 45% at SNR =2 for the Amplitude, FHDM, delay, and TTP. The TransientAnalyzer performed much better for the Amplitude and FDHM; it performed worse in the case of TTP and delay estimation. However, unlike the TransientAnalyzer, the software SpikeAnalyzer (3) cannot be used for transients of different origins. The software CalTrack for calcium transients analysis (9) was not recommended by the authors for transients with SNR less than 15. The GPR approach thus provides to TransientAnalyzer the precision and accuracy better than or comparable to the specialized image-based or function-based approaches and bears the advantage of general applicability.

The finite precision of the parameter estimates opens the question of the experimental design needed to reach the required confidence in the estimated parameter values. Assuming a normal distribution of estimates, the number of replications necessary for a certain relative precision can be calculated from the coefficient of variation *C_V_* (32). For instance, for the mean of *N* determinations to be within 5% of the true value with a 95% probability, *N* = 4 is sufficient if *C_V_* = 5% but *N* = 67 is needed if *C_V_* = 25%.

Similarly, to detect a 20% effect of an experimental intervention on a parameter at the significance level of p < 0.05 with 95% power, one would need to compare 4 transients before and 4 transients after the intervention if *C_V_* = 5%, 16 transients before and 16 transients after the intervention if *C_V_* = 15%, but 42 transients before and 42 transients after the intervention if *C_V_* = 25%.

### Capabilities

The TransientAnalyzer software provides a general solution to the common problem of the exact characterization of noisy transient signals. First, it corrects the baseline, detects the transients, and finds their onset time in a continuous record. Second, it approximates the detected transients one by one with a Gaussian process regression and creates the corresponding spline function. The output is a noise-free continuous function that allows an exact estimation of characteristic parameters of individual transients at a high amplitude and time resolution. In this way, also the response of the properties of the transients to experimental interventions can be analyzed. The application is independent of the device and method of detection. As demonstrated above, TransientAnalyzer reliably estimates the characteristic parameter values of transient changes in cellular calcium concentration. In the same way, it could be used to estimate parameters of cell shortening occurring in cultured or isolated myocytes, or various optical, electrical, mechanical, or chemical signals related to specific cell activities.

The examples of detection and approximation of transient signals of various origins are given in Figure 5. The results of the approximation by a corresponding mathematical model are compared with the approximation by the model-free GPR method (panels A - D). In these randomly selected examples, the GPR method provided similar or smaller residuals than the model-fitting methods. In the case when the experimental trace did not contain artifacts and the model was a good description of the trace, the two methods provided equivalent mean squared error (Figure 5 A, B). Otherwise, the mean squared error of the model-fitting method was up to twice as large as the error of the GPR method estimate (Figure 5 C, D). Nevertheless, for transients that can be described by a mechanistic (not phenomenological) model function, a model-fitting approach may allow more exact interpretations. Algorithms based on model fitting may faithfully analyze even data too noisy for estimation by model-free algorithms (8). On the other hand, the GPR-based approach may be especially useful when the process that generates the transient is complex and not well understood. In such cases, reliable estimation of transient characteristics recorded in various experimental situations may help to discover the underlying mechanisms.

**Figure 5.**
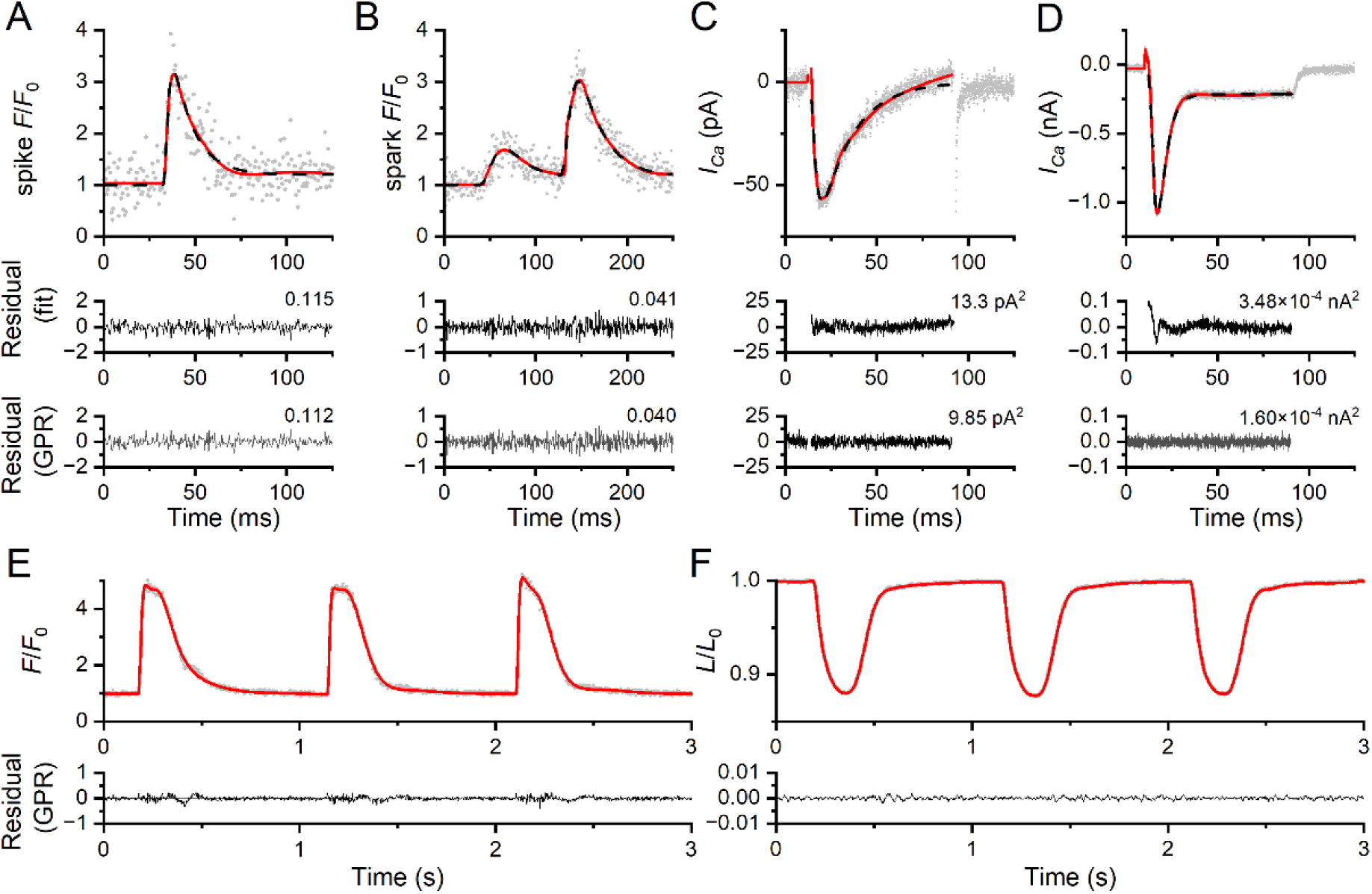
Examples of analysis of transients by GPR. A - a calcium spike recorded in a rat cardiac myocyte in response to stimulation (11); B - two sequential calcium sparks recorded in a rat cardiac myocyte (11); C - calcium current recorded in a neonatal rat cardiac myocyte in response to stimulation (36); D - calcium current recorded in an adult rat cardiac myocyte in response to stimulation (37); E, F - calcium transients and sarcomere length transients recorded in a rat cardiac myocyte in response to 1-Hz field stimulation (38). Points - the recorded signals. Red lines - results of analysis by the GPR method, black dashed lines - results of fitting with mathematical models used in the respective cited studies. The bottom panels show residuals between the experimental data and their approximation by GPR. The mean squared errors are given next to the residuals.

For convenience, the application is published at (https://github.com/IuliiaBaglaeva/TransientAnalyzer) as the Python package TransientAnalyzer, which can be installed by the generally available Package Installer for Python (33) using the command *pip install TransientAnalyzer*, and as a graphical user interface created using PyQt5 (34) and pyqtgraph (35). The core module of the TransientAnalyzer documented in (https://transientanalyzer.readthedocs.io/en/latest/) allows customization of the software by the user for a specific purpose.

### Limitations

The detection part is based on the maximal difference between the recorded and filtered signals. It can be expected that large instances of noise could result in false detection of a transient or incorrect determination of its onset. On the other hand, small and slowly rising transients may be missed. This problem can be resolved by the trial-and-error method using the interactive design of the GUI.

It should be noted that the activation phase was the critical part of the tested signal. In the simulated transients with a rise time of *t*_10-90_% of 12.5 ms, the rising phase was sampled by only five data points. Despite this, the approximation software based on the GPR method performed robustly. Nevertheless, the high and low-frequency noise may cause substantial errors when the amplitude of the noise is comparable with that of the signal, as discussed in the Results section. From this point of view, the number of outliers was acceptable. The functionalities of TransientAnalyzer were not tested for peak SNR smaller than 2. It should be also noted that the reported errors of measurement are solely due to the normally distributed noise in the signal records. Experimental conditions that warrant stable transient signal records and normal distribution of noise can be sometimes difficult to meet in the real world. However, the comparison of the expected and observed precision can help to optimize the experimental design by identifying the sources of excessive signal variability.

The GPR method is computationally intensive. The complexity caused by matrix inversion increases with the 3^rd^ power of the number of data points. This may result in a somewhat prolonged analysis in the case of transients with a large number of data points. For example, the 100 simulated calcium transients analyzed in the Results section (400 data points per transient) took approximately 100 seconds on a PC with 6 core (12 threads) Intel® Core™ i5-11400H. However, the GPR module uses GPFlow based on TensorFlow technology, which may accelerate substantially the learning and prediction of GPR by the use of a dedicated GPU. For example, the analysis of the same 100 simulated calcium transients took approximately 50 seconds on a PC with Intel® Core™ i5-11400H and NVIDIA® GeForce RTX™ 3060 Laptop GPU. Alternatively, downsampling of records or omission of the steady part of the baseline can be used if individual transients contain more than 500 points to speed up calculations on less powerful computers. Since the algorithm does not require regular sampling periods, non-constant sampling periods can be used during signal acquisition or post-recording, if necessary.

## Conclusions

The TransientAnalyzer is a self-standing computational tool designed for the automatic analysis of transient signals in records with a high noise level that provides reliable estimates of the characteristic parameters of detected transients. The software TransientAnalyzer including the core module is documented to allow its extension and customization for specific applications by a knowledgeable user of the Python system.

## Supporting information

Supplement: User Guide

## Author Contributions

IB, AZ, IZ, and BI designed research; IB and BI performed research; IB and BI contributed analytic tools; IB and AZ analyzed data; IB, BI, AZ, and IZ wrote the paper.

## Declaration of Interest

The authors declare no competing interests.

## Funding

The research was supported by the SAV-TUBITAK project JRP/2019/836/RyRinHeart, by the project VEGA 2/0182/21, the Slovak Research and Development Agency project APVV-21-0443, the ERA-NET project ERACVD_JTC2019-055/HF-MetaB to A. Zahradníková, Jr.,and by the Operational Programme Integrated Infrastructure for the projects: Long-term strategic research of prevention, intervention, and mechanisms of obesity and its comorbidities, IMTS: 313011V344, and Strengthening of research, development and innovation capacities of translational biomedical research of human diseases, IMTS: 313021BZC9, co-financed by the European Regional Development Fund.

## Notes

https://github.com/IuliiaBaglaeva/TransientAnalyzer

https://github.com/IuliiaBaglaeva/TransientAnalyzerUI

https://github.com/IuliiaBaglaeva/TransientAnalyzerUI/releases

### Competing Interest Statement

The authors have declared no competing interest.

### Summary of Updates

Added another source of funding Added a footnote that the manuscript has been accepted to the Biophysical Journal under doi:10.1016/j.bpj.2023.01.003.

