## Supplement: User Guide for "Analysis of noisy transient signals based on Gaussian process regression"

### Transient Analyzer
### User Guide

#### Introduction

TransientAnalyzer is a software for analysis of the time course of transient records that allows automatic detection of transients in the record, their best approximation by the Gaussian process, the construction of a surrogate spline function, and the estimation of signal parameters, including the peak amplitude, onset time, time to peak, full duration at half maximum, rise and decay times, and transient durations. The TransientAnalyzer UI can be run on a PC with Windows 7 and higher.

Python libraries PyQt5, GPFlow, Pyqtgraph, NumPy, SciPy, and Pandas are used in this software.

The software TransientAnalyzer including its core module is documented in https://github.com/IuliiaBaglaeva/TransientAnalyzer, which allows its extension and customization for specific applications. Knowledge of the Python system is required.

To use TransientAnalyzerUIEXE.zip, download it from https://github.com/IuliiaBaglaeva/TransientAnalyzerUI/releases/tag/Release. This archive contains files TransientAnalyzer.exe and TransientAnalyzer.ui. Unzip this archive into one folder.

The data for testing are in the directory https://github.com/IuliiaBaglaeva/TransientAnalyzerUI/tree/main/Examples.

#### Chapter 1. User interface and data input .

Open the TransientAnalyzer.exe file. The screen depicted in Figure 1 will appear.

The main Program window contains on the left side from top to bottom: the Menu bar with tabs **File** and **Help**, the Infobar, the Graph panel, and the Input panel. On the right side, there is the Table panel for viewing the results of the analysis. The Progress bar is at the bottom of the window.

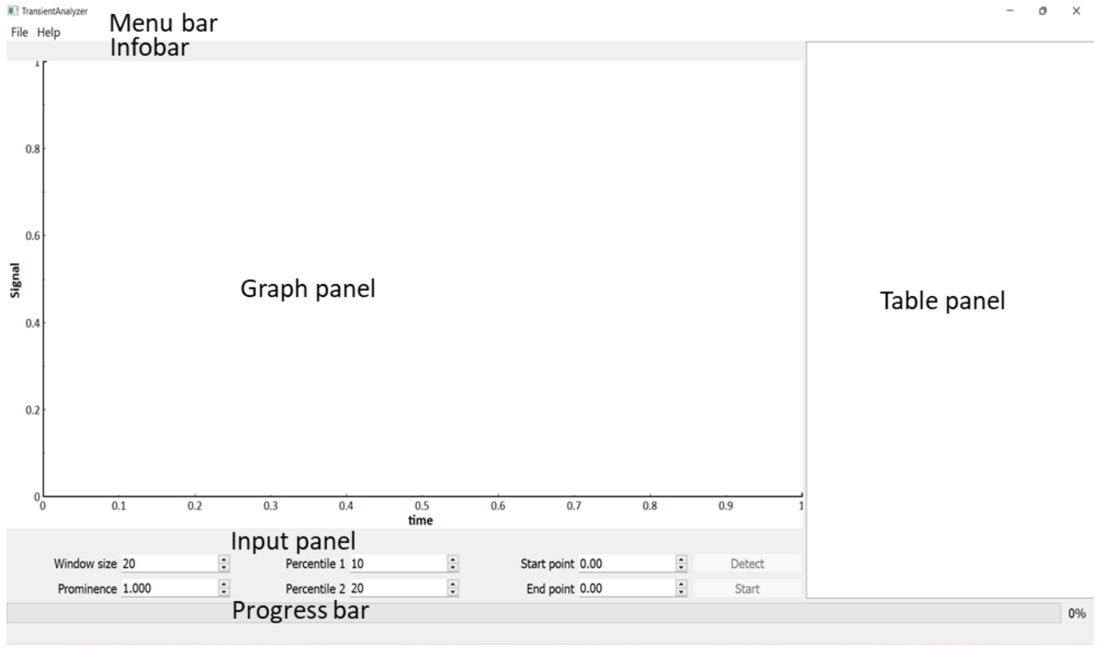

Figure 1. The main window of the GUI.

The content of the tab **File** (Figure 2):

- **Open File** - selection and opening of the data file for analysis. The allowed file formats are .txt, .csv or .xlsx. The .txt file data should be space-separated. The first column should contain the independent variable of the recorded trace, i.e., sequential number, time value, position, etc. The second column should contain values of the measured signal (fluorescence, position, voltage, current, power, etc.). For data in .xlsx and .csv formats, the first row must contain the names of columns. If the names of columns are written in the format "**name, [unit]",** the program uses the unit in the output table of the estimated parameter values; otherwise, the parameters in the table will be in arbitrary units.
- **Add stimulation File -** selection and opening of the optional stimuli data file specifying the time points of stimuli applied during signal recording. It should have either a single column with exact values of stimulation times or two columns with the independent variable in the first column and "0" or "1" in the second column to indicate when the stimulus occurred. The number of rows has to be the same as the length of the corresponding signal record data file.
- **Save results -** saves the table of the estimated parameter values of the detected transients and the whole approximated trace in full resolution. The Results file is saved in .xls format that contains two sheets - "Parameters" and "Traces".
- **Exit**

The tab **Help** contains information about this product.

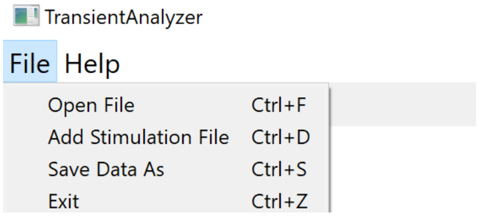

**Figure 2. The File menu**

The Input panel (Figure 3) contains text boxes with a set of parameters controlling the analysis of transients with default values that can be modified by the user to tune the performance and modify the output of the analysis:

The box **"Window size"** allows setting the number of points used for the boxcar filter (the default value is 20). The size of the boxcar filter affects the estimation of the transient starting time value. Depending on sampling frequency and transient kinetics, changing the window size may improve the accuracy of *the start time* estimation (see Chapter 3: Troubleshooting).

The box **"Prominence"** allows changing the threshold value for the minimal peak amplitude of a transient that will be automatically detected and approximated by the GPR kernel (the default value is 1). When the recorded signal has a low signal-to-noise ratio, decreasing the Prominence may cause increased detection of false transients, and vice versa (see Chapter 3: Troubleshooting).

The boxes **"Percentile 1"** and **"Percentile 2"** specify two amplitude levels, defined as percentage of the peak amplitude, used for calculations of kinetic characteristics of the transient. For instance, the **Percentile** 1 value of 10 specifies the calculations of the rise time as the time difference between 10 and 90 % ($t_{10\text{-}90\%}$) of the peak amplitude. At the same time, the **Percentile** 2 value of 20 specifies the calculation of the rise time as the time difference between 20 and 80 % ($t_{20\text{-}80\%}$) of the peak amplitude. These percentile values also specify calculations of the transient duration from the start to the time of 90 % or 80 % decrease from the peak amplitude ($d_{90\%}$, $d_{80\%}$), respectively.

The boxes **"Start point"** and **"End point"** allow setting the range of the analyzed data record (see below).

In the upper left corner of the Input panel, the x and y values of the cursor position on the displayed trace are shown.

The Progress bar at the bottom of the Program window (Figure 1) indicates the progress of the analysis in %.

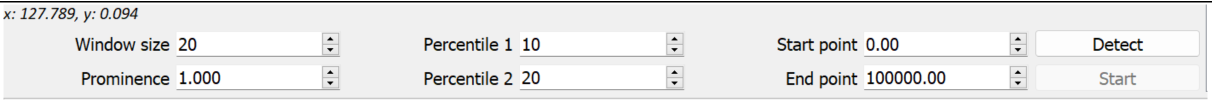

**Figure 3. The Input panel**

### Chapter 2. Analysis and data output.

Open the tab **File,** click the **Open File** menu item, and select the file for analysis. In the Graph panel, the whole data set is displayed (Figure 4). The name and the directory path of the input file are displayed in the Infobar. In the case of transients recorded in response to stimulation, the option **Add stimulation File** allows the selection and opening of the stimulation data file. The software recognizes the stimulation data file in .txt, .csv, or .xlsx formats. Uploading of the stimulation file is reported in the Infobar.

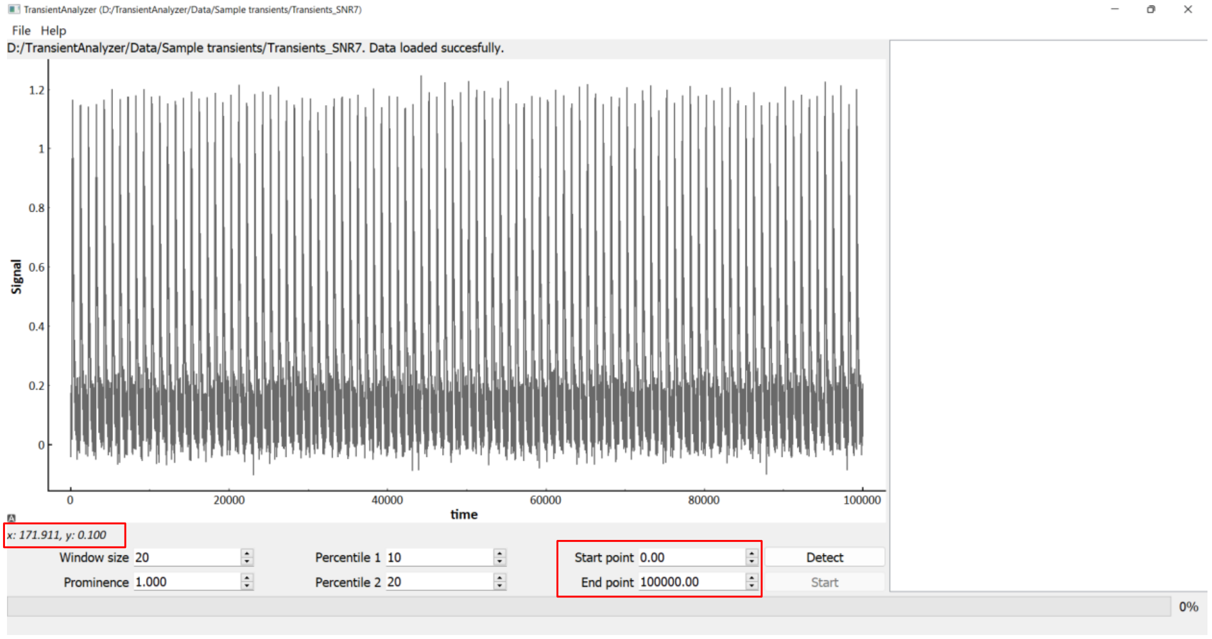

**Figure 4. The data file upload**

After uploading the data file, the region of the recorded trace intended for analysis can be selected. Use the mouse zooming and panning tool described in Table 1 to inspect the recorded data. By default, the whole trace is selected and displayed in the Graph panel, which is indicated by the time values of the first and the last point of the trace in the Start and End point boxes of the Input panel (Figure 4). To inspect the recorded signal, use the mouse zooming and panning tools described in Table 1.

The segment of the trace for analysis  can be determined easily. Click into the Start point box, then position the cursor to the first data point trace to be displayed, and double-click on it. This will change the value accordingly in the Start point box. Alternatively, the required x value for the Start point of analysis can be directly written in the box. The last point can be specified in the same way using the End point box. The specified trace segment will be displayed in the Graph pannel.

**Table 1. Control of the graphical display by the mouse**
(https://pyqtgraph.readthedocs.io/en/latest/mouse_interaction.html):

| Action | Mouse control |
|---|---|
| Zoom in the y-axis | Right button + Moving up |

| | |
|---|---|
| Zoom out the y-axis | Right button + Moving down |
| Zoom in the x-axis | Right button + Moving right |
| Zoom out the x-axis | Right button + Moving left |
| Zoom in and zoom out | Wheel moving |
| Graph panning | Left button |

Finally, the Percentile values for estimation of the decay and rise times are specified (Figure 5). The default values of 10 and 20 correspond to the 10% - 90% and 20% - 80% percentiles).

Click the **Detect** button to launch the detection of transients. The progress is indicated at the bottom of the screen. After a few seconds, the Start time points of all detected transients are indicated by vertical dashed lines. Use the mouse zooming and panning tools described in Table 1 for inspection of details. The total number of detected transients is displayed in the Infobar (Figure 5).

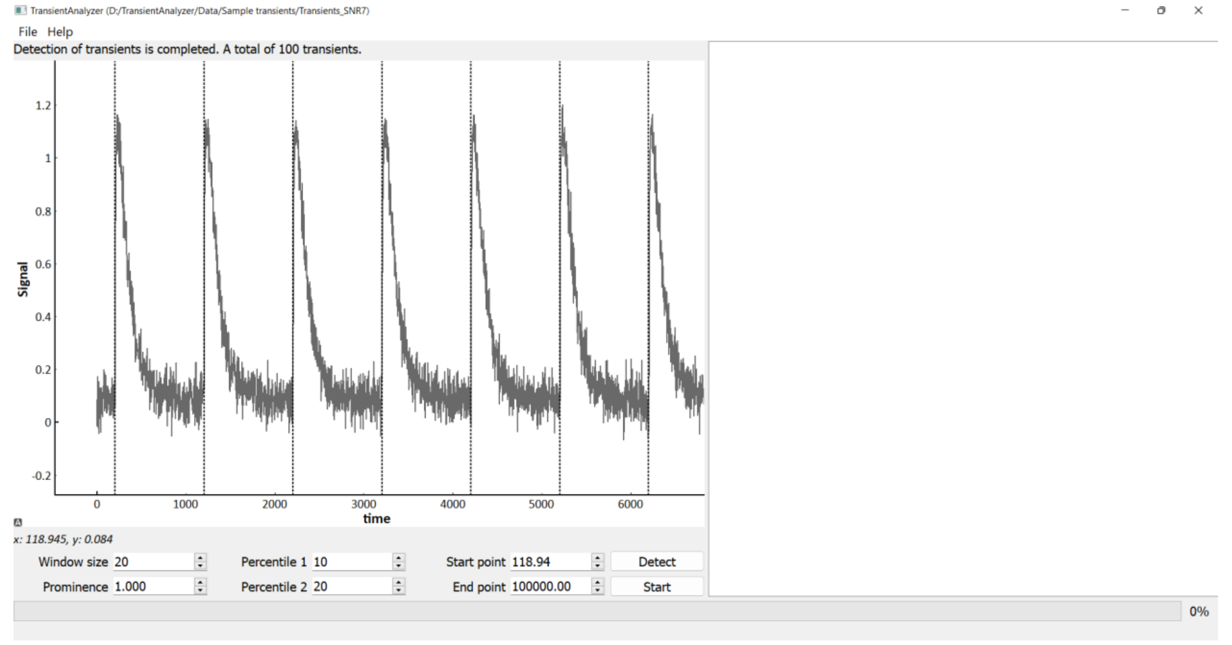

**Figure 5. Detection of the transients**

When all transients were correctly detected or the file with the stimulation times was uploaded successfully, the GPR analysis of transients can be activated by clicking the **Start** button. The progress bar displays the progress of the analysis. The Infobar shows the message “Parameter estimation in progress”. When the parameter estimation process is complete, the spline of the best approximation curve (red) is overlaid above the analyzed signal record and the Results table containing the estimated parameter values of all detected transients is displayed on the right side of the Program window (Figure 6). The divider between the graph and the table windows can be moved by the mouse.

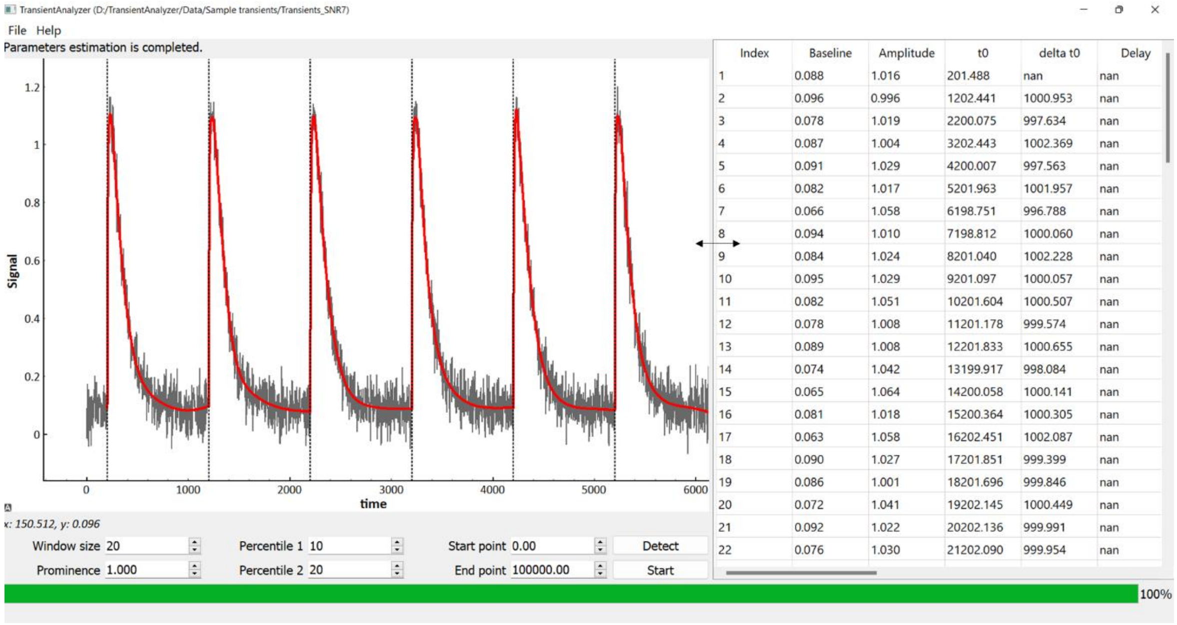

**Figure 6. The results of parameter estimation.**

The parameters in the Results table defined in Figure 7 are:

- **Index** – the sequential number of the transient.
- **Baseline** – the value of the baseline
- **Amplitude** – the peak amplitude
- **t_0** – $t_0$, the start time value of the transient.
- **Delta t_0** – the mean interval between transients.
- **Delay** – the difference between the stimulus and $t_0$
- **t_10 - 90 %** and **t_20 - 80 %** are the rise times for 10% to 90% and 20% to 80% of the peak amplitude, respectively.
- **TTP** – *TTP*, the time to peak
- **FDHM** – *FDHM*, the full duration at half-maximum (FDHM)
- **t_90 - 10 % and t_80 – 20 %** are the decay times between 90% to 10% and 80% to 20% of the peak amplitude.
- **d_80%** and **d_90%** are the durations from the start of the transient to the time of 80% and 90% decrease from the peak amplitude, respectively.

If the input data files contained units, they are included in the table heading. If the input data did not contain the units (i.e., if the input was in the .txt format or if the user did not provide units of the data columns, the output looks like the example in Figure 7):

- The units of Baseline and Amplitude values correspond to that of the y-axis
- The units of the remaining parameter values correspond to that of the x-axis.

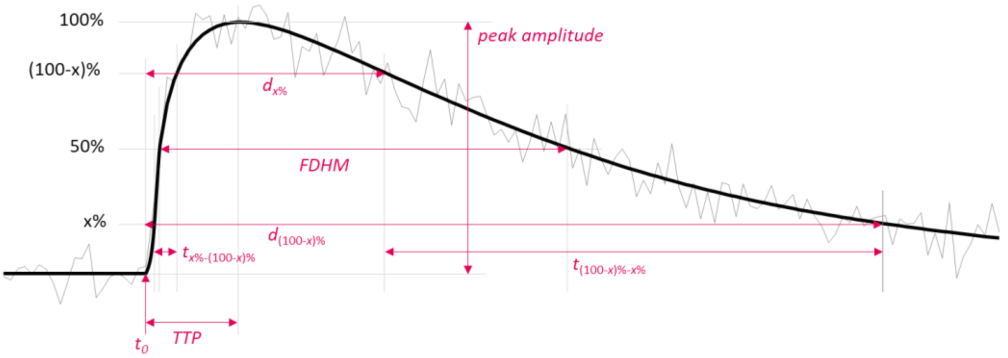

TransientAnalyzer (D:/TransientAnalyzer/Data/Sample transients/Transients_SNR7)

File Help

| Index | Baseline | Amplitude | t0 | delta t0 | Delay | t_10%-90% | t_20%-80% | TTP | FDHM | t_90%-10% | t_80%-20% | d_90% | d_80% |
|---|---|---|---|---|---|---|---|---|---|---|---|---|---|
| 1 | 0.088 | 1.016 | 201.488 | nan | nan | 11.258 | 6.411 | 28.510 | 141.130 | 274.090 | 172.628 | 335.761 | 252.896 |
| 2 | 0.096 | 0.996 | 1202.441 | 1000.953 | nan | 9.369 | 5.860 | 34.762 | 146.212 | 251.956 | 156.625 | 319.514 | 243.530 |
| 3 | 0.078 | 1.019 | 2200.075 | 997.634 | nan | 12.817 | 8.134 | 35.732 | 145.596 | 276.147 | 176.195 | 343.680 | 263.546 |
| 4 | 0.087 | 1.004 | 3202.443 | 1002.369 | nan | 11.867 | 7.044 | 34.086 | 142.944 | 274.243 | 168.712 | 344.208 | 255.762 |
| 5 | 0.091 | 1.029 | 4200.007 | 997.563 | nan | 15.674 | 10.363 | 30.875 | 142.213 | 272.295 | 177.753 | 330.501 | 256.360 |
| 6 | 0.082 | 1.017 | 5201.963 | 1001.957 | nan | 11.374 | 6.571 | 26.617 | 145.019 | 268.851 | 167.880 | 336.028 | 253.788 |
| 7 | 0.066 | 1.058 | 6198.751 | 996.788 | nan | 13.031 | 8.187 | 32.509 | 143.438 | 305.761 | 186.330 | 365.146 | 265.694 |
| 8 | 0.094 | 1.010 | 7198.812 | 1000.060 | nan | 14.403 | 8.755 | 36.238 | 142.122 | 268.231 | 164.135 | 336.636 | 253.031 |
| 9 | 0.084 | 1.024 | 8201.040 | 1002.228 | nan | 13.670 | 8.994 | 29.457 | 140.616 | 279.497 | 174.865 | 341.834 | 257.579 |
| 10 | 0.095 | 1.029 | 9201.097 | 1000.057 | nan | 16.694 | 10.017 | 32.421 | 142.911 | 265.048 | 168.662 | 324.833 | 248.115 |

**Figure 7. Characteristics of a transient.** Top – an example of a simulated transient (noisy gray line) with its best approximation (smooth solid line) and the graphical definition of the characteristic transient parameters. Bottom – an example of the estimated parameter values as displayed in the Result table.

### Chapter 3. Troubleshooting.

Errors in the detection of transients caused by the noise can be fixed by a proper setting of the "Window size" and "Prominence" parameters.

In the case of missed transients, it is recommended to decrease the value **of Prominence,** as shown in Figure 8.

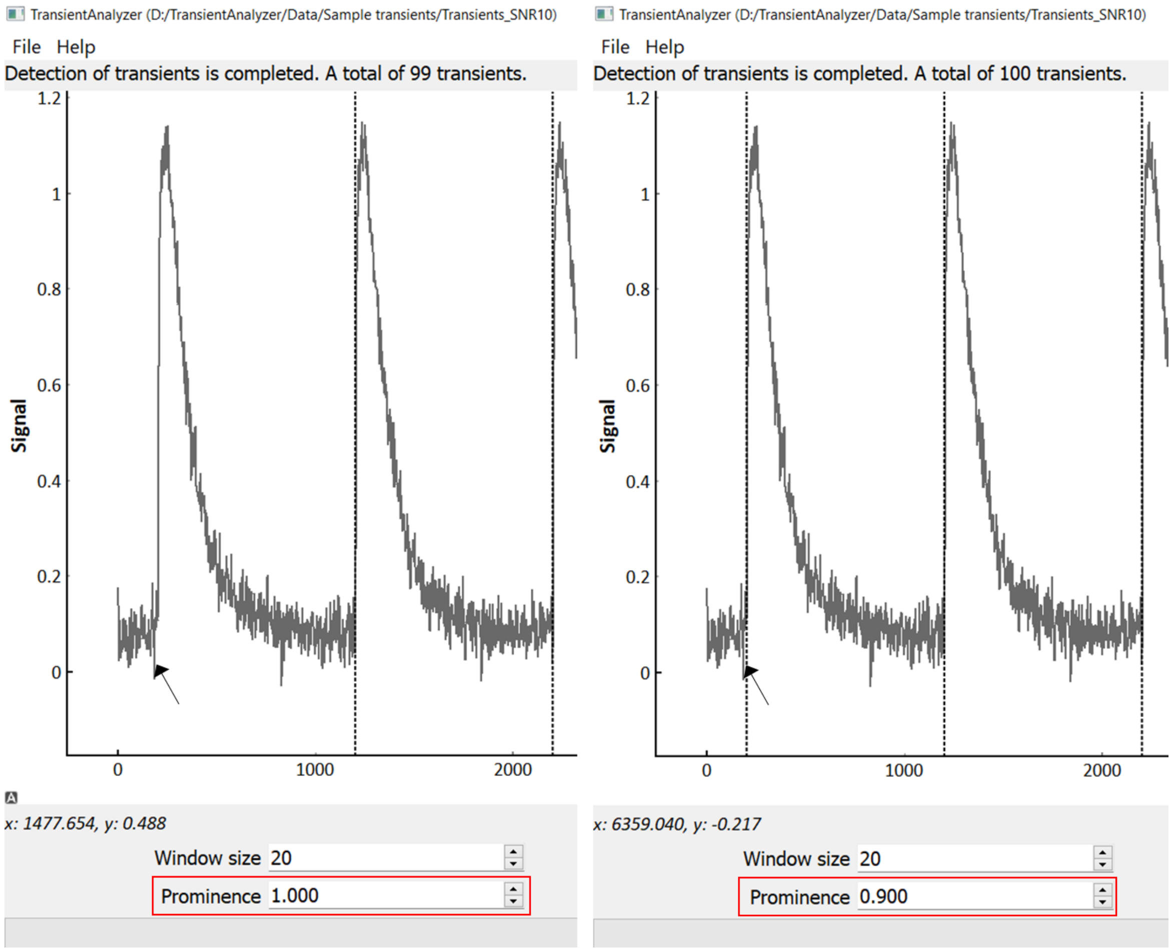

Figure 8. Problem with missing transient detection. Left: the missed transient (arrow) at the default value of **Prominence.** Right: correct detection. The problem was solved by decreasing **Prominence** to 0.9.

In the case of false detection of transients, the **Window size** should be increased first (Figure 9). If false transients persist, the **Prominence** should be increased as well.

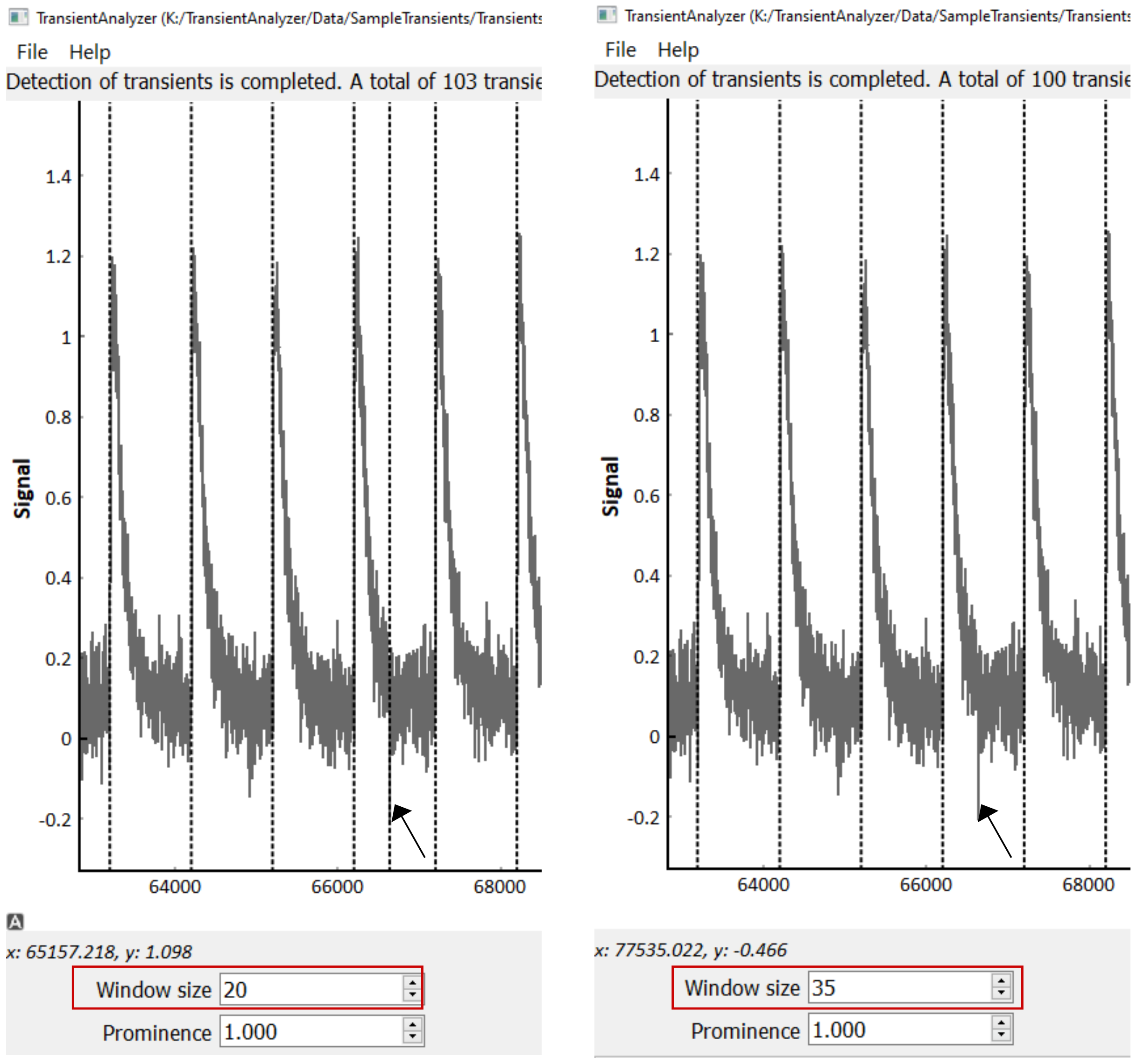

Figure 9. Problems with false transient detection. Left - a false transient detected at the default Window size. Right - the problem was solved by increasing Window size to 35.
